# In Vitro Expansion of Keratinocytes on Human Dermal Fibroblast-Derived Matrix Retains Their Stem-Like Characteristics

**DOI:** 10.1101/381178

**Authors:** Chee-Wai Wong, Beverley F. Kinnear, Radoslaw M. Sobota, Rajkumar Ramalingam, Catherine F. LeGrand, Danielle E. Dye, Michael Raghunath, E. Birgitte Lane, Deirdre R. Coombe

**Affiliations:** School of Pharmacy and Biomedical Sciences, Faculty of Health Sciences, Curtin University, Bentley, WA 6102, Australia; Curtin Health Innovation Research Institute, Faculty of Health Science, Curtin University, Bentley, WA 6102, Australia; Institute of Molecular and Cell Biology, Agency for Science, Technology and Research (A*STAR), 61 Biopolis Drive, No. 07-48A Proteos, Singapore 138673, Singapore; Institute of Medical Biology, Agency for Science, Technology and Research (A*STAR), 8A Biomedical Grove, No 06-06 Immunos, Singapore 138648, Singapore; Centre for Cell Biology and Tissue Engineering, Competence Centre for Tissue Engineering and Substance Testing (TEDD), Institute for Chemistry and Biotechnology, ZHAW School of Life Science and Facility Management, Zurich University of Applied Science, Switzerland; Centre for Cell Therapy and Regenerative Medicine, School of Biomedical Sciences, The University of Western Australia, Crawley, WA

**Keywords:** Extracellular matrix, keratinocytes, adult stem cells, serum-free culture, xenogeneic-free culture, self-renewal

## Abstract

The long-term expansion of keratinocytes under serum- and feeder free conditions generally results in diminished proliferation and an increased commitment to terminal differentiation. Here we present a serum and xenogeneic feeder free culture system that retains the self-renewal capacity of primary human keratinocytes. *In vivo*, the tissue microenvironment is a major contributor to determining cell fate and a key component of the microenvironment is the extracellular matrix (ECM). Accordingly, acellular ECMs derived from human dermal fibroblasts, cultured under macromolecular crowding conditions to facilitate matrix deposition and organisation, were used as the basis for a xenogeneic-free keratinocyte expansion protocol. A phospholipase A_2_ decellularisation procedure produced matrices which, by proteomics analysis, resembled in composition the core matrix proteins of skin dermis. On these ECMs keratinocytes proliferated rapidly, retained their small size, expressed p63, did not express keratin 10 and rarely expressed keratin 16. Moreover, the colony forming efficiency of keratinocytes cultured on these acellular matrices was markedly enhanced. Collectively these data indicate that the dermal fibroblast-derived matrices support the *in vitro* expansion of keratinocytes that maintained stem-like characteristics under serum free conditions.

## Introduction

The skin is an indispensable barrier that safeguards the body from the external environment. It possesses the ability to self-renew, which enables the replacement of dead cells and the repair of wounds thereby sustaining a barrier function[1]. In normal circumstances, most cutaneous wounds heal without medical intervention. However, if the wound is extensive and extends into the dermis, medical attention may be required[2]. Traditionally, the therapeutic strategy for treating large, deep wounds has been to use split-thickness skin autografts. However, this treatment is not viable in the case of extensive burn injury, as patients may lack sufficient healthy donor sites[3].

The grafting of cultured keratinocytes is an alternative treatment to assist in the repair of damaged skin. This method uses a technique originally developed by Rheinwald and Green[4] to expand keratinocytes *in vitro* from a patient’s skin biopsy. In this method, expansion of keratinocytes is achieved using an irradiated mouse fibroblast feeder layer and medium containing foetal bovine serum (FBS). While this is effective for rapidly expanding keratinocytes, the reliance on xenogeneic components carries a potential risk of exposing patients to animal pathogens and immunogenic molecules[5]. To address these concerns, *in vitro* culture systems that omit both the feeder layer and serum were developed. A popular system uses a defined serum-free medium that contains the necessary growth factors and a collagen matrix to support keratinocyte attachment and growth[6, 7].

While this defined culture system may meet regulatory approval, its ability to propagate keratinocytes is inferior to the Rheinwald and Green[4] system. Keratinocytes grown in the defined serum-free system have a more limited lifespan; a diminished self renewal capacity and an increased commitment towards differentiation or senescence[7, 8]. This suggests that the defined serum-free system does not fully meet keratinocyte requirements. It is likely crucial elements required to sustain undifferentiated keratinocytes long term, reside in the fibroblast feeders used in the Rheindwald and Green system. Fibroblasts secrete cytokines, growth factors and extracellular matrix (ECM). The focus for defined culture systems has been on the cytokines and growth factors[9, 10], but it is possible the ECM is a crucial requirement that has been overlooked.

The ECM is complex meshwork of macromolecules, comprising fibrous structural proteins (e.g. collagen, fibronectin, laminin and elastin), specialised proteins (e.g. growth factors) and proteoglycans (e.g. perlecan). It was previously thought to be an inert structure that provided a platform for cell adhesion, but it is now known that the ECM provides both biochemical and biomechanical cues that regulate cell behaviours such as adhesion, migration, proliferation and differentiation[11, 12]. Currently, there is considerable interest in using cell-derived matrices to reproduce a tissue specific microenvironment. Numerous studies have shown that acellular ECM assists in maintaining the stem cell phenotype and in promoting self-renewal during *in vitro* expansion[13–16]. However, keratinocyte expansion on a dermal fibroblast derived-matrix (Fib-Mat) under serum free conditions has not been well examined.

While it is possible to generate an acellular ECM *in vitro*, most culture methods produce an unstructured ECM that lacks critical components[17, 18]. This may be due, in part to differences between the *in vitro* and *in vivo* microenvironments. Cells in culture are in a dilute solution of macromolecules of 1-10 mg/ml, which is several-fold lower than the normal physiological environment that can range from 20.6 to 80 mg/ml[19]. Thus, in culture, molecular interactions in the extracellular environment may not be sufficient to produce an ECM which resembles that seen *in vivo*. To mitigate this problem, the addition of large, inert macromolecules such as Ficoll™ to the culture medium has been used to mimic the density of macromolecules within tissues. Molecules like Ficoll™, when used in this context, have been called “macromolecular crowders” (MMC) and the process of mimicking the *in vivo* concentration of macromolecules is called “macromolecular crowding”. Interestingly, the addition of Ficoll™ to cell cultures was found to accelerate biochemical reactions and supramolecular assembly, and macromolecular crowding was found to affect the deposition and architecture of the ECM[17, 18, 20].

Here, we describe the development and functional characterization of a xenogeneic-free matrix derived from primary human dermal fibroblasts (Fig 1). A proteomics analysis confirmed that this matrix resembled, in its core protein composition, the ECM of human dermal tissue. When used as a substrate for keratinocyte growth in the absence of feeder cells and under defined serum-free conditions this Fib-Mat facilitated keratinocyte proliferation. In addition, more keratinocytes maintained the stem-like characteristics of small cell size, expression of p63 and a lack of keratin 16 expression as well as the retention of a colony forming capability. These data indicated that these acellular Fib-Mat are an appropriate microenvironment to enable the expansion of undifferentiated keratinocytes *in vitro*.

**Figure 1:**
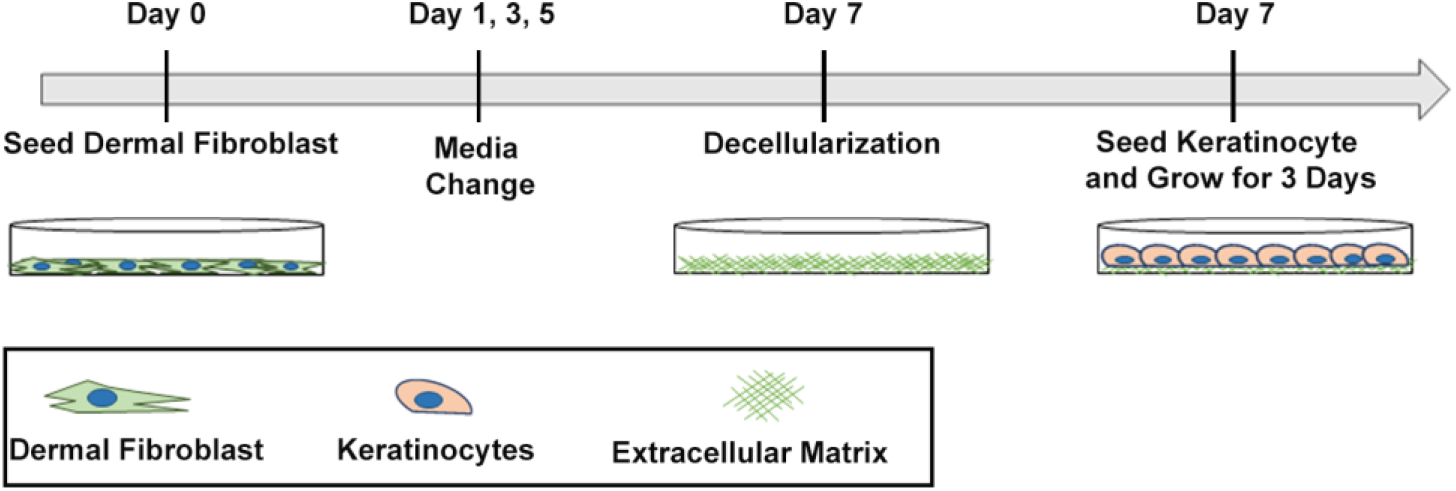
Schematic of the preparation of a xenogeneic-free acellular dermal fibroblast-derived matrix as a substrate for keratinocytes.

## Materials and Methods

### Antibodies

The primary rabbit polyclonal antibodies used were anti-type I collagen (Abcam; Cambridge, UK), anti-type IV collagen (Abcam), anti-fibronectin (Abcam) and the anti-perlecan antibody CCN-1was a gift from Prof. John Whitelock (University of NSW, Sydney, Australia). The mouse monoclonal antibodies (mAbs) used were: anti-fibroblast marker (clone TE7; Millipore; MA, USA), anti-involucrin (clone SY5; Sigma; MO, USA), anti-ki67 (clone MM1; Novacastra; Wetzlar, Germant), anti-p63 (clone 4A4; Abcam), anti-α-smooth muscle actin (clone 1A4; Sigma), anti-Thy1 (BD Bioscience; NJ, USA), and anti-vimentin (clone V9; Dako; CA, USA). The following mouse mAbs: anti-keratin 10 (K10, clone LH2), anti-keratin 14 (K14, clone LL001) and anti-keratin 16 (K16, clone LL025) were produced in house. The secondary antibodies used were Alexa488 anti-mouse IgG, Alexa546 anti-mouse IgG, Alexa 488 anti-rabbit IgG and Alexa546 anti-rabbit IgG (all from Molecular Probes, ThermoFisher Scientific; OR, USA).

### Cell cultures

Two human dermal fibroblast (HDF) donors were used, one cell population was obtained from the American Type Culture Collection (ATCC; VA, USA) and the other was from the Skin Cell Bank of Institute of Medical Biology, Singapore, and the use of these cells was covered by ethical codes overseen by Curtin University Human Ethics committee (ethics approval number: HRE-2016-0273A). HDFs were maintained in Dulbecco’s Modified Eagle’s medium (DMEM) supplemented with 10% FBS (Serana Europe GmBH; Pessin Germany), 10 mM HEPES, 2 mM L-glutamine and 1 mM sodium pyruvate (Gibco. ThermoFisher Scientific). Human neonatal keratinocytes (purchased from Gibco) were cultured on tissue culture growth surfaces that have been coated with type I collagen (Sigma) in PBS (3μg/cm^2^) and maintained in Defined Keratinocyte Serum Free medium (DKSFM; Gibco). Cells were maintained in tissue culture incubators at 37 °C and 5 % carbon dioxide. Keratinocytes at passage 4 or 5 were used for all experiments.

### Extracellular matrix deposition with macromolecular crowding treatment

HDFs were seeded at a density of 15,000 cells/cm^2^ and were allowed to attach overnight in basal medium comprising DMEM: Ham F12 (3:1) supplemented with 2% human serum (Gibco), 10 mM HEPES, 2 mM L-glutamine, 1 mM sodium pyruvate and 30μg/ml ascorbic acid (Wako Chemical; Tokyo, Japan). The medium was then replaced with fresh medium containing 7.5 mg/ml Ficoll 70 (Sigma) and 25 mg/ml Ficoll 400 (GE Lifesciences; Buckinghamshire, UK) to induce macromolecular crowding. The HDFs were cultured for 6 days for ECM deposition, with the medium changed on alternate days.

### Decellularization of dermal fibroblast-derived extracellular matrix

The ECMs deposited using macromolecular crowding were decellularised using either EDTA (ET), ammonium hydroxide (AH) or phospholipase A_2_ (PLA_2_). For the ET method, cells were rinsed with PBS followed by 2.5 mM EDTA/PBS, and then incubated in 2.5 mM EDTA/PBS for 10 min at 37°C. Using a P1000 pipet, the cell monolayer was sprayed off leaving the matrix. The matrix was washed with PBS, incubated for 5 min at 37°C with 0.5% Triton X-100/PBS and washed with PBS. For AH decellularisation the cells were washed with PBS and incubated in 0.02 M ammonium hydroxide (Sigma)/0.5% Triton X-100/1x EDTA-Free protease inhibitor (Roche; Basel, Switzerland) at 37°C for 5 min. For the PLA_2_ method the cells were washed in PBS and incubated in PLA_2_ (20 U/ml) (Sigma)/50 mM Tris-HCl (pH 8)/0.15 M NaCl/1 mM MgCl_2_/1 mM CaCl_2_/0.5% sodium deoxycholate/1x EDTA-Free protease inhibitor (Roche) at 37°C for 30 min. Matrices decellularised by the AH and PLA_2_ methods were then washed with PBS before being treated with 0.02 mg/ml DNase I (Amresco, PA, USA) in reaction buffer (10 mM Tris-HCl (pH 7)/2.5 mM MgCl_2_/0.5 mM CaCl_2_) at 37°C for 30 min and then washed again with PBS. The presence of DNA in decellularized ECM was determined by staining with 4′,6-diamidino-2-phenylindole (DAPI; Sigma; 1μg/ml in PBS), while the presence of residual actin was determined by staining with 1 unit/ml of phalloidin conjugated with tetramethylrhodamine (TRITC). Images were captured using a Zeiss LSM510 inverted fluorescent microscope. In addition to visualizing DNA using DAPI, residual nucleic acids in the decellularized ECM were measured using the CyQUANT cell proliferation assay kit (Molecular Probes, Thermo Fisher Scientific), following the manufacturer’s protocol. Fluorescence intensity was measured with 485 nm/535 nm filters using an EnSpire Multimode Plate Reader (Perkin Elmer; MA, USA).

### Immunocytochemistry analysis

Immunofluorescent staining was performed on cells or ECM adhered to etched glass coverslips in 24-well plates. Coverslips were prepared as described in Chaturvedi *et al*[21]. The cell or ECM layer was fixed with 4% paraformaldehyde in PBS for 15 min at RT, washed with PBS then blocked with 10% Goat Serum/1% BSA/PBS for 1h at RT. Blocking solution was removed, and samples were incubated for 1 h at RT with primary antibody prepared in 10% Goat Serum/1% BSA/PBS. Cells were washed 3 x 5 min with PBS before being incubated for 1 h with secondary antibodies prepared in 10% Goat Serum/1% BSA/PBS. The ECMs were washed 3 x 5 min with PBS and incubated in DAPI (1μg/ml in PBS) for 10 min. Coverslips were mounted in Vectashield antifade mounting medium (Vector Laboratories; Peterborough, UK) and sealed with nail varnish. Images were captured with either a Zeiss Axioskop Fluorescent Microscope (Carl Zeiss, Germany) using Spot Advanced software (Michigan, USA) or a Nikon A1+ Confocal Microscope (Nikon, Tokyo, Japan). All antibodies were titrated to determine their appropriate concentration for the experiment. To generate a 3D representation of the matrix, Z-stacked images of anti-Type I Collagen antibody stained ECM were obtained using a Nikon A1+ Confocal Microscope and images were merged using the NIS-Elements AR analysis software.

### Keratinocyte Proliferation

The proliferation of keratinocytes on the substrates (Fib-Mat, type I collagen (3 μg/cm^2^) and tissue culture plastic (TCP) was assessed. Keratinocytes were harvested and seeded at a density of 1 x 10^4^ cells/well in a 48-well tissue culture plate (NUNC, ThermoFisher Scientific) and grown for six days. At 24 h intervals keratinocytes were fixed for 5 min with cold acetone:methanol (1:1), washed with PBS and incubated with PBS/1%BSA for 1 h at RT before the nuclei were stained with DAPI (Sigma). Using an Olympus IX-81 high content screening inverted microscope (Olympus; Tokyo, Japan) and a 10x objective, 7 by 11 non-overlapping quadrants were imaged, to produce a 0.5 cm^2^ area image. Nuclei/cell numbers were determined using Fiji-Image J software and its “Find Object” macro.

### Keratinocyte Adhesion to Substrates

Cell adhesion assays were performed in 96 well tissue culture plate (NUNC). Keratinocytes were harvested, resuspended in adhesion assay buffer (DMEM (Gibco), 10 mM HEPES (Gibco), 2 mM L-glutamine (Gibco), 1 mM sodium pyruvate (Gibco), 0.2% BSA (Sigma), 25 μg/ml adenine (Sigma), 0.4 μg/ml hydrocortisone (SOLU-CORTEF^®^, Pfizer; NY, USA), 0.12 IU/ml insulin (Humulin^®^, Lilly; IN, USA), and seeded at a density of 1 x 10^4^ cells/well and left to adhere for 1 h at 37°C to either decellularized HDF ECM, type I collagen (3 μg/cm^2^) or TCP. Unbound cells were removed by washing with adhesion assay buffer followed by PBS. The plate was placed overnight at −80°C and brought up to RT before cell number was determined using the CyQUANT cell proliferation assay kit (Molecular Probes, Invitrogen), following manufacturer’s instructions. Fluorescence intensity was measured with 485nm/535nm filter using an EnSpire Multimode Plate Reader (Perkin Elmer). Results were calculated as a percent of the control, which was prepared by pelleting 1 x 10^4^ keratinocytes, washing and storing the pellet at −80°C before using the CyQuant assay kit to determine the fluorescence intensity of this number of keratinocytes.

### Keratinocyte Size and Motility

Keratinocytes were harvested and seeded (1 x 10^4^ cells/well) on the various substrates in the wells of a 48-well culture plate (NUNC). After three days of culture in DKSFM keratinocytes were fixed with 4% paraformaldehyde/PBS for 15 min at RT, incubated in PBS/1%BSA for 1 h at RT before polymerized actin was stained with 1 unit/ml of Phalloidin-Alexa 488 (Molecular Probe) and nuclei with DAPI. Using an Olympus IX-81 high content screening inverted microscope (Olympus), 8 by 8 non-overlapping quadrants were imaged using a 20x objective lens. Cell size as delineated by the stained actin cytoskeleton was determined using the Cell Profiler software.

Keratinocyte movement was assessed on either Fib-Mat, type I collagen coating or uncoated TCP. Keratinocytes were seeded as described and following an overnight incubation to allow cell adhesion, live images were taken using an Olympus IX-81 high content screening inverted microscope (Olympus). Time-lapse images were taken at 15-min intervals over 2 days using a 10x objective.

### Quantification of Ki67 Positive Keratinocytes

Briefly, keratinocytes were harvested and then seeded as described above and at day 3, keratinocytes were fixed with 4% paraformaldehyde for 15 min at RT, permeabilized with cold 0.1% Triton X-100/PBS for 3 min and then incubated in PBS/1% BSA for 1 h at RT. The keratinocytes were blocked with 10% Goat Serum/1% BSA/PBS for 1 h at RT before being incubated with a 0.3 μg/ml anti-Ki67 antibody (Novacastra). This was followed by a 1 h incubation with Alexa 488 conjugated anti-mouse antibody at RT. Keratinocyte nuclei were stained using DAPI. Images were taken on the Olympus IX-81 high content screening inverted microscope. Using a 10x objective, 7 by 11 non-overlapping quadrants were imaged. The percent of keratinocytes positive for Ki67 was determined using the Cell Profiler software.

### Quantification of p63 Positive Keratinocytes

Keratinocytes were seeded onto the different substrates and cultured for 3 days after which they were fixed, permeabilised and blocked as described for the Ki67 expression experiment. Keratinocytes were then incubated with 0.5μg/ml of anti-p63 mAb and the anti-mouse Alexa 546 conjugated antibody, washed and the slides mounted in a DAPI-containing anti-fade mounting medium. A Nikon A1+ confocal microscope and a 10x objective lens was used to image 3 by 3 non-overlapping quadrants per slide. The percent of p63 positive keratinocytes was determined using Cell Profiler software.

### Colony Forming Efficiency Assay

Keratinocytes were grown on acellular HDF-derived ECM, type I collagen coating or uncoated TCP for 3 days in DKSFM, before being harvested. Keratinocytes (1 x 10^3^) from each substrate were seeded into 6-well tissue culture plates containing mitomycin-treated 3T3-J2 feeder cells and cultured for 10-12 days in culture medium which was a 3:1 ratio of DMEM (Gibco) and Ham’s F12 (Gibco) supplemented with 10 mM HEPES (Gibco), 2 mM L-glutamine (Gibco), 1 mM sodium pyruvate (Gibco), 25 μg/ml adenine (Sigma), 0.4 μg/ml hydrocortisone (SOLU-CORTEF^®^, Pfizer), 0.12 IU/ml insulin (Humulin^®^), 2 nM triiodothyronine (Sigma), 10 ng/ml epidermal growth factor (BD Bioscience) and 5 mM forskolin (Sigma). Media changes were performed on alternate days. After 10-12 days the cells were fixed with (1:1) acetone: methanol and stained with 0.1% toluidine blue in ddH2O. The colonies that formed were counted, with colonies ≥ 1 mm^2^ being “large” and the rest “small”.

### Mass Spectrometry and Proteomics Analysis

Dermal fibroblast-derived ECMs were generated using macromolecular crowding and decellularized using PLA_2_. To solubilize the acellular ECM, 8 M Urea/50 mM Tris-HCl pH 8.0 was added before scraping the matrix off the surface and transferring it to a microtube. The matrix mixture was reduced with 10 mM DTT (Sigma), alkylated with 55 mM Iodoacetamide (Sigma) and then diluted with 100 mM TEAB buffer to reach a Urea concentration of < 1 M. The matrix proteins were digested with sequencing grade endoproteinase Lys-C (Promega, WI, USA) and sequencing grade-modified trypsin (Promega) at a ratio of 1:100 at 25^0^ C for 4 h and 18 h respectively, samples were subsequently acidify with 1% TFA and desalted. Following desalting with a Sep-Pak C18 column cartridge (Waters, Milford MA), the samples were analysed using an Easy nLC 1000 liquid chromatography system (Thermo Fisher Scientific) coupled to an Orbitrap Fusion Mass Spectrometer (Thermo Fisher Scientific). Each sample was analyzed in a 60 min gradient using an Easy Spray Reverse Phase Column (50 cm x 75 μm internal diameter, C-18, 2 μm particles, Thermo Fisher Scientific). Data were acquired in −3 s cycle with the following parameters: MS in Orbitrap and MS/MS in ion trap with ion targets and resolutions (OT-MS 4 x E5 ions, resolution 120 K, IT-MS/MS 1000 ions/turbo scan, “Universal Method”).

Data analysis: The peak list was generated using Proteome Discoverer (Version 1.4. Thermo Fisher Scientific). The MS/MS spectra were searched with the Mascot 2.5.1 (Matrix Science, MA, USA) search algorithm using the Human UniProt Database. The following parameters were used: precursor mass tolerance (MS) 20 ppm, IT-MS/MS 0.6 Da, 3 missed cleavages; Variable modifications: Oxidation (M), Deamidated (NQ), Acetyl N-terminal protein, Static modifications: Carboamidomethyl (C). Forward/decoy search was used for false discovery rate (FDR) estimations on peptide/PSM level, and were set at high confidence, FDR 1% and medium confidence, FDR 5%. The generated protein list was curated using the Matrisome[22] database.

### Statistical Analyses

Statistical analyses were performed using SPSS statistics software V22.0 (IBM Corporation, NY). If the data sets were normally distributed and their variances homogeneous, a parametric test was conducted; if not, a non-parametric test was used. A p-value of p≤0.05 was considered statistically significant. For normally distributed data, experiments containing two treatments data sets were analysed with a t-test. For experiments containing 3 or more treatment data sets, one-way analysis of variance (ANOVA) was conducted followed by Tukey’s posthoc test. As p63 expression data were not normally distributed, these were analysed using the Kruskal-Wallis one way analysis of variance followed by Wilcoxon’s signed-ranked test.

## Results

### Development of an Acellular Dermal Fibroblast-Derived Matrix

To produce a matrix that best mimics the microenvironment of keratinocytes would encounter *in vivo*, primary human dermal fibroblasts (HDFs) from adult donors were chosen as the cell source. Phase contrast microscopy revealed that the HDFs used had a uniform spindle-like morphology that is typical of fibroblasts. Immunofluorescence analyses indicated they expressed the fibroblast markers TE-7, thy-1 and vimentin (Fig. S1).

It was reported that the addition of a mixture of Ficoll 70 and Ficoll 400 to the culture medium has the effect of mimicking the space occupied by glycoproteins in plasma, and this was found to benefit ECM deposition by cells *in* vitro[17]. Accordingly, the Ficoll cocktail was included in the fibroblast culture medium to act as MMC. The Ficoll cocktail marginally altered the appearance of the HDFs, and reduced their proliferation over a 7 day period (Fig. S2A, C). However, absent immunoreactivity with an α-SMA antibody indicated the HDFs had not differentiated into myofibroblasts. In contrast, when the Ficoll cocktail was ommitted and the cells were cultured for 7 days α-SMA staining was detected, and cell morphology of the α-SMA positive cells resembled that of myofibroblasts (Fig. S2B). Immunostaining of the ECM deposited when HDFs were cultured with the Ficoll MMC for 7 days revealed an increase in the staining intensity of type I collagen, type IV collagen, fibronectin and perlecan compared to cultures without MMC (Fig. 2A). When visualized with confocal microscopy, a layer of type I collagen was seen fully embedding HDFs in MMC culture, also covering the top of the cells, whereas cultures without MMC had markedly less type I collagen staining (Fig. S3A). Removal of the HDFs using a phospholipase A_2_ (PLA_2_) decellularisation technique revealed that type I collagen and fibronectin were deposited uniformly across the surface when HDFs were cultured with MMC, when compared to cultures without MMC (Fig. S3B).

**Figure 2:**
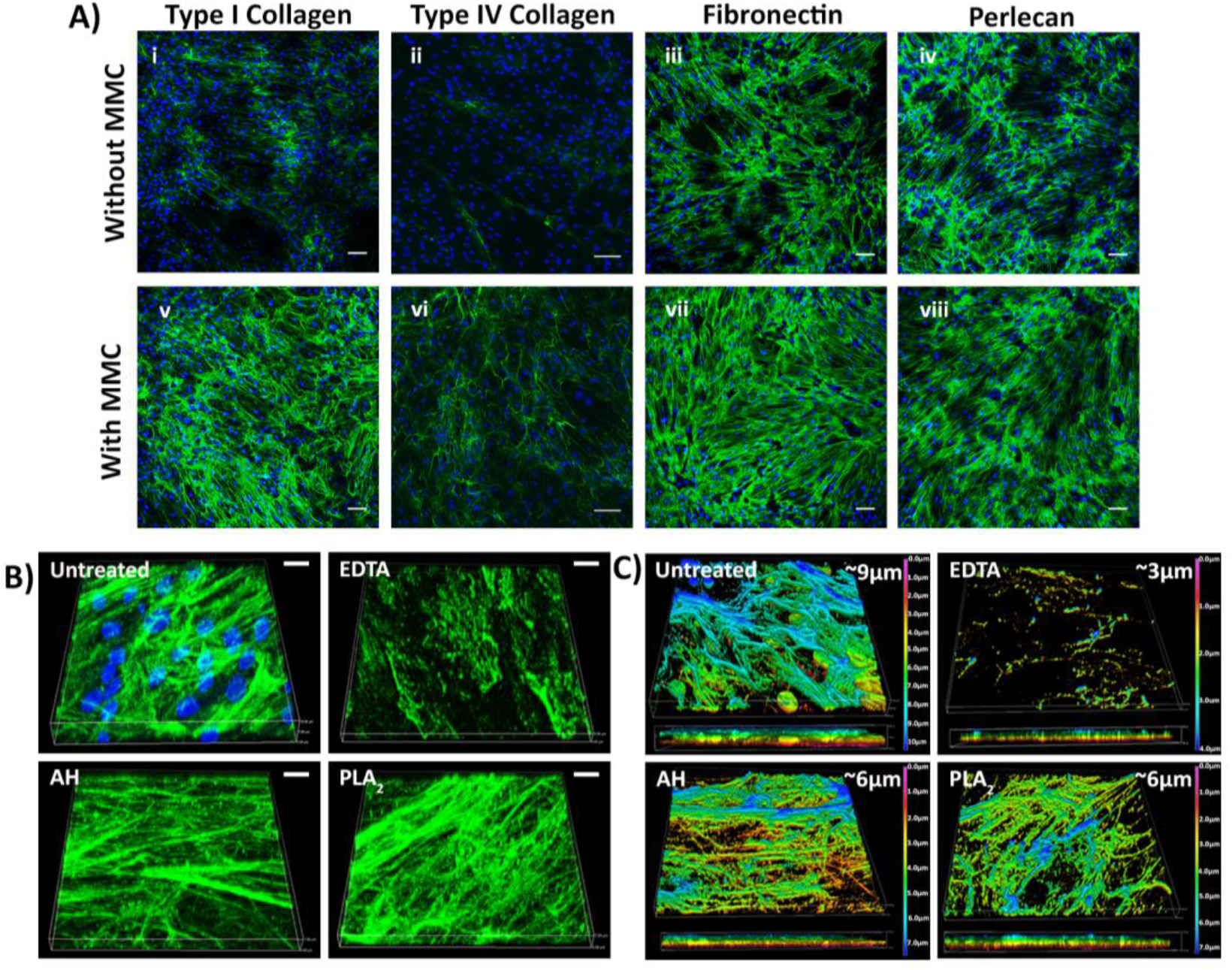
ECM proteins deposited by dermal fibroblasts. **A)** The effect of MMC on HDF ECM deposition. The cells and matrix were immunostained for type I collagen (i, v), type IV collagen (ii, vi), fibronectin (iii, vii) and perlecan (iv, viii). Scale bars are 100 μm. **B)** The 3D architecture of the ECM deposited by HDF before and after decellularisation with EDTA, AH or PLA_2_ as revealed by type I collagen immunostaining and DAPI staining. Scale bars are 10 μm. **C)** The thickness of the ECM after decellularisation. The colour coding represents the Z-depth location within the 3D z-stacked image.

As macromolecular crowding enabled a well-structured and uniform deposition of ECM, without myofibroblast differentiation, MMC were used to generate the Fib-Mat. To determine a decellularisation method which removed the cells, yet preserved the ECM proteins and structure as much as possible, three decellularisation protocols were compared. These were: EDTA, ammonium hydroxide (AH) and PLA_2_. Phase contrast microscopy revealed all three protocols removed the fibroblasts. However, fibril-like structures were retained only with AH and PLA_2_ treatments (Fig. S4). DAPI staining of the nucleic acids remaining after decellularisation indicated both the EDTA and the PLA_2_ methods more effectively removed DNA than the AH method, which left distinct nuclear fragments in the ECM (Fig. S5Ai). Quantification of DNA removal indicated that the EDTA and PLA_2_ methods were effective in removing 99% of the DNA, whereas 97% of the DNA was removed with the AH method (Fig. S5Aii). To determine whether the cytoskeletal components of the HDFs were removed following decellularisation, phalloidin-TRITC staining was used to detect actin filaments. As shown in Figure S5B, a few actin filament fragments were detected following AH treatment, but no staining was apparent when EDTA or PLA_2_ treatments were used.

To investigate the structure of the ECM following decellularisation, 3D Z-stacked confocal images were obtained following type I collagen immunostaining. Fibrillar structures of type I collagen resembling the non-decellularised control were clearly visible following AH and PLA_2_ treatments, but the EDTA treatment disrupted the structure of the type I collagen filaments (Fig. 2B). The thickness of the ECM was measured by determining the depth of type I collagen staining was 9 μm thick. After decellularisation using the AH or PLA_2_ protocols, ECM thickness decreased to around 6 μm, which dropped further to 3 μm following EDTA treatment (Fig. 2C). Immunofluorescence intensities of the variously decellularised matrices revealed a significant reduction in type I collagen and fibronectin staining after EDTA treatment. In contrast, both AH and PLA_2_ treatments were shown to preserve type I collagen and fibronectin immunostaining (Fig. S6).

### Dermal Fibroblast Matrices Mimics Skin Dermis Extracellular Matrix

Collectively the data demonstrated that the PLA_2_ decellularisation protocol produced an intact ECM that was devoid of most cell components, hence this method was used to generate acellular matrices (Fib-Mat) for further analyses. As the goal was the production of an acellular matrix that mimicked the dermal ECM, the protein compositions of the ECM derived *in vitro* from two dermal fibroblast donors were determined using mass spectrometry (MS)-based proteomics. To investigate whether the ECM produced *in vitro* by dermal fibroblasts matched dermal ECM, the ECM signature of the dermis was obtained from the “Human Protein Atlas” database[23] and was compared to our proteomic data. However, after examination of the “Human Protein Atlas”, it was apparent that a number of core ECM proteins like collagen III alpha 1 (COL3A1) and laminin alpha-4 (LAMA4), which are present in the dermis, were not found in the “Human Protein Atlas” database. To ensure a comprehensive list of dermal ECM proteins was used for the comparison, the proteomic dataset of the skin prepared from studies by Bliss *et al*.[24] was used to supplement the “Human Protein Atlas” database. To curate the ECM proteins, the proteomic analyses and the “Human Protein Atlas” supplemented data were categorised using the human matrisome database (MatrisomeDB, http://matrisomeproject.mit.edu/). This database categories the ECM proteins into “core matrisome” (ECM glycoproteins, collagens and proteoglycans) and “matrisome-associated proteins” (ECM-affiliated proteins, ECM regulators and secreted factors)[22]. This analysis revealed that most of the core matrisome proteins expressed in the skin dermis were also found in the Fib-Mat that were prepared (Fig. 3). However, the majority of the matrisome-associated proteins were absent in these Fib-Mat (Fig. 3).

**Figure 3:**
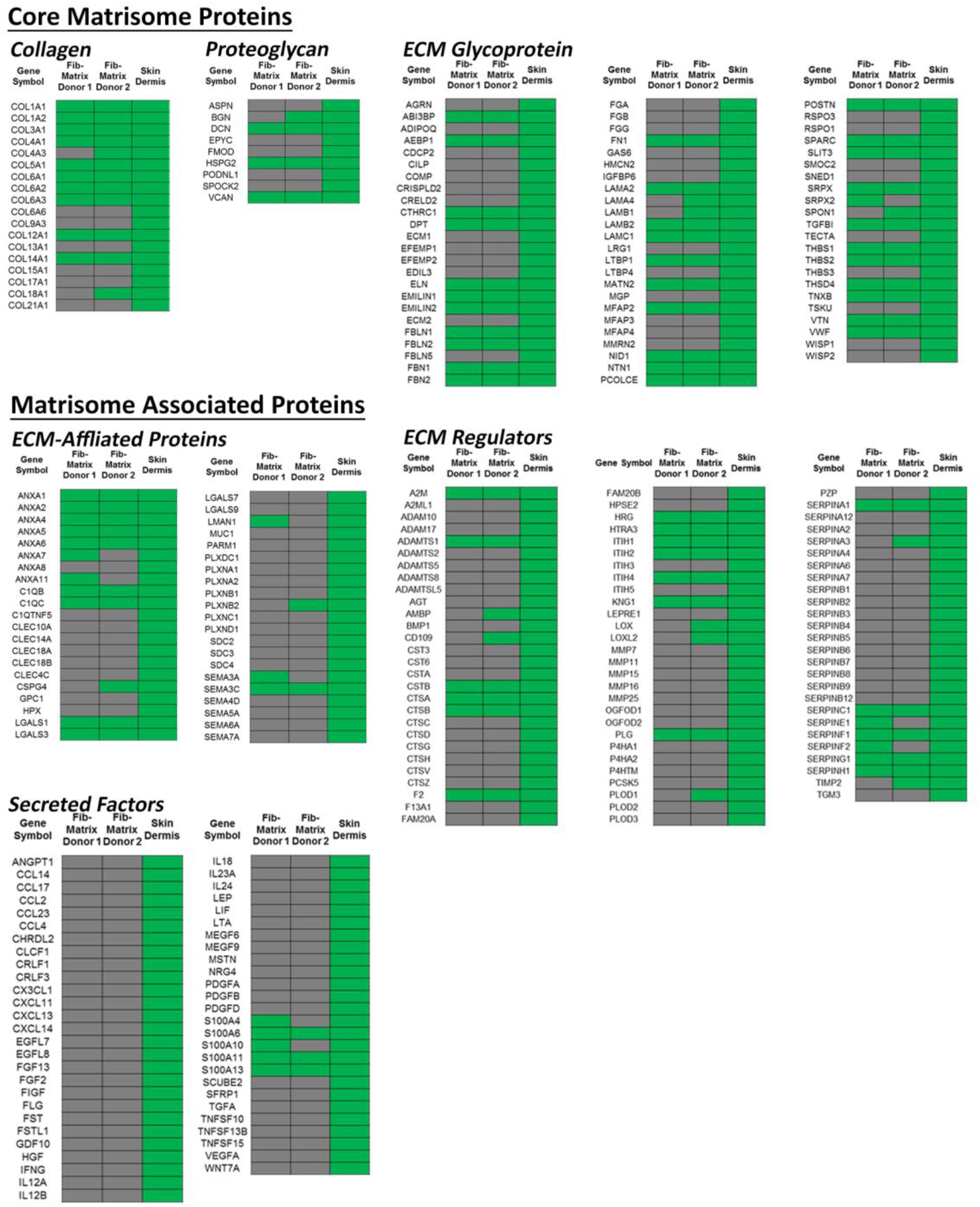
Protein composition of the acellular ECM. Protein compositions of the acellular ECM from dermal fibroblasts from two donors were compared to that of the dermis. The ECM proteins are subdivided into two categories; “core matrisome” (ECM glycoproteins, collagens and proteoglycans); and “matrisome-associated proteins” (ECM-affiliated proteins, ECM regulators and secreted factors). Green and grey indicate the presence or absence of ECM proteins respectively.

### Dermal Fibroblast-Derived Matrix Supports Keratinocyte Proliferation

Next, the ability of Fib-Mat to support the adhesion and proliferation of keratinocytes was investigated. Type I collagen was used as a positive control, as this is the substrate commonly used with defined keratinocyte serum free medium (DKSFM) for propagating keratinocytes[6, 7]. Tissue culture plastic (TCP) was the negative control. The extent of keratinocyte adhesion to the different substrates differed. More keratinocytes attached to Fib-Mat (84%), than to type I collagen (77%) or TCP (56%; Fig. 4A). Phase contrast microscopy revealed that keratinocytes adhered well to both Fib-Mat and type I collagen but less to TCP. While keratinocytes proliferated on all three substrates, their behaviour differed. On Fib-Mat keratinocytes grew as colonies, and cells within the colonies had a small cobblestone morphology, which persisted until day 6 whereupon a near confluent monolayer was reached. Although similar behaviour was observed on TCP, the keratinocytes comprised a heterogeneous population of differing sizes. Whereas, keratinocytes on type I collagen grew as single cells, they also have a mixed population, with some of the cells being large and flat (Fig. 4B & Fig. 6A).

**Figure 4:**
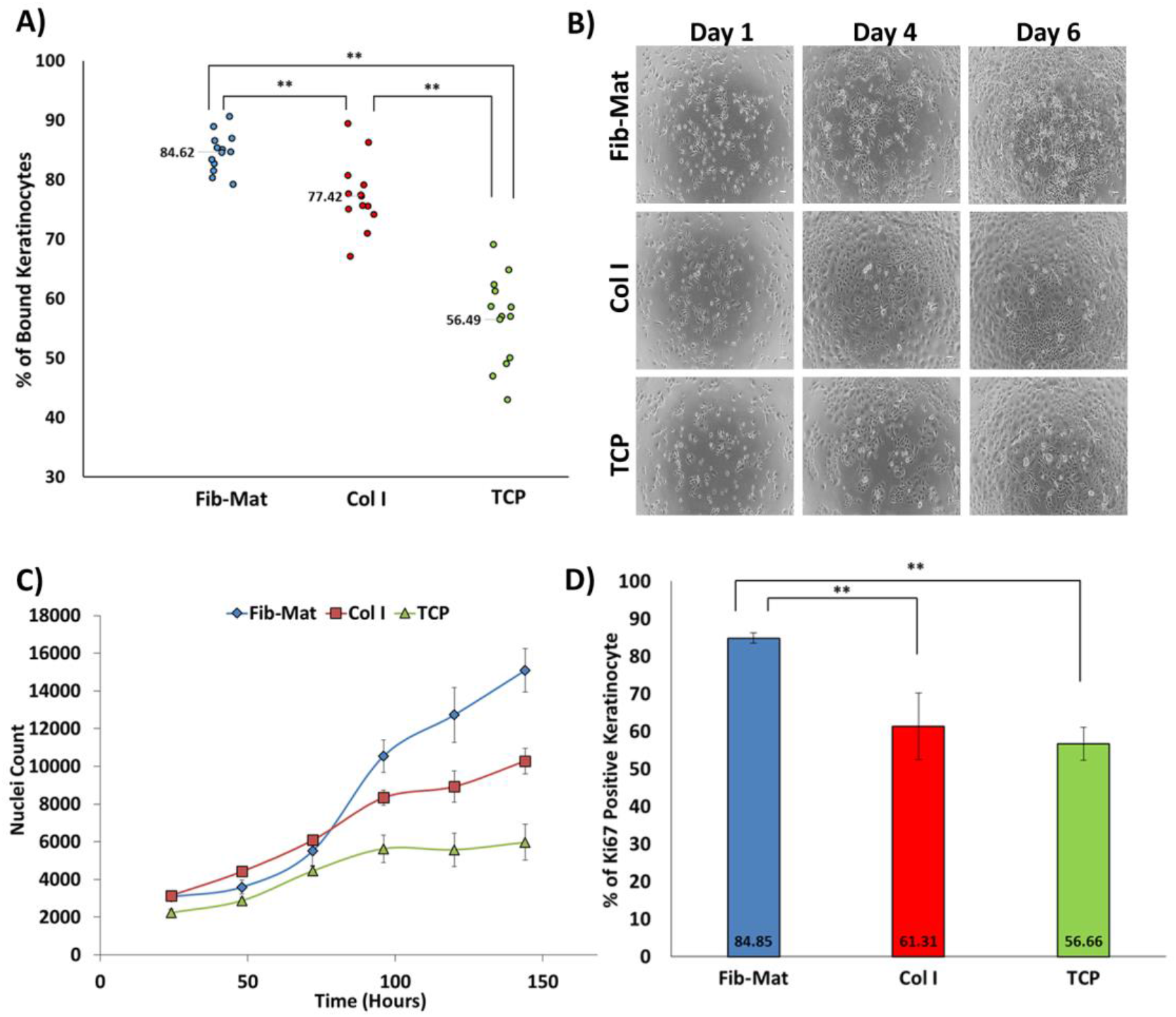
The different substrates support keratinocyte adhesion and proliferation to varying degrees. **A)** The ability of Fib-Mat, Col I and TCP to support keratinocyte adhesion. The data are the mean percent of bound keratinocytes ± standard deviation; mean values are given. Shown are pooled data from three separate experiments. **= P<0.01 **B)** Morphology of keratinocytes growing on dermal fibroblast-derived matrix (Fib-Mat), type I collagen (Col I) and tissue culture plastic (TCP) as captured by phase contrast microscopy. Keratinocyte on days 1, 4 and 6 post seeding are shown. Scale bars are 100 μm. **C)** The ability of Fib-Mat, Col I and TCP to support keratinocyte proliferation. Nuclei were stained with DAPI and counted. The data are the mean ± standard deviation obtained from 4 replicate wells. A representative of three separate experiments is shown. **D)** Ki67 expression by keratinocytes cultured on Fib-Mat, Col I and TCP. The data are the mean percent of Ki67 positive keratinocytes ± standard deviation; mean values are given. The data shown are representative of three separate experiments. ** = P<0.01.

The rate of keratinocyte proliferation on the different substrates also differed. On Fib-Mat keratinocytes initially proliferated more slowly than keratinocytes on type I collagen but reached similar numbers by day 3. Thereafter, an exponential rate of proliferation was observed in keratinocytes on Fib-Mat, which was higher than that seen on type I collagen. On TCP, keratinocyte proliferation was slow and the rate plateaued by day 4 (Fig. 4C). Keratinocyte expression of Ki67 was determined on day 3, as the growth curve (Fig. 4C) indicated a change in proliferation rates on each of the substrates at this point. More keratinocytes on Fib-Mat (84.85%) stained with the Ki67 mAb compared to that seen in keratinocytes on type I collagen (66.31%) and TCP (56.66%; Fig. 4D). Determination of the numbers of Ki67 expressing cells on day 4 and 5 revealed this remained the case (data not shown). From these data Fib-Mat was the best substrate to promote keratinocyte proliferation.

### Keratinocytes Grown on Dermal Fibroblast-Derived Matrix were Undifferentiated

To determine differentiation status, keratinocytes grown on the different substrates were immunoststained for K16, K14, K10, and involucrin. To acquire better image resolution at high magnification, etched glass coverslips (EGC) were used. To check if this growth surface affected keratinocyte behaviour, keratinocytes were grown on either TCP or EGC coated with Fib-Mat or type I collagen, or were uncoated. Cells were grown for 3 days before being fixed and immunostained for K14. Keratinocytes grown on all surfaces were similarly positive for K14 (Fig. S7). On EGC coated with type I collagen, keratinocytes grew as colonies, whereas they grew as single cells on TCP coated with type I collagen. As this was the only change, immunostaining of the differentiation markers was performed on keratinocytes grown on EGC with, or without, the various coatings. Although K14 expression was observed in keratinocytes on all substrates a population of keratinocytes on uncoated EGC did not express K14 (Fig. 5A). While the expression of K10 was not observed in keratinocytes on any substrate, involucrin expression was detected in cells on all substrates. However, a higher proportion of keratinocytes were positive for involucrin when grown on uncoated EGC, as compared to keratinocytes on Fib-Mat or type I collagen. In addition, A higher proportion of keratinocytes grown on type I collagen and EGC compared to keratinocytes grown on Fib-Mat were observed to express K16 (Fig. 5A).

**Figure 5:**
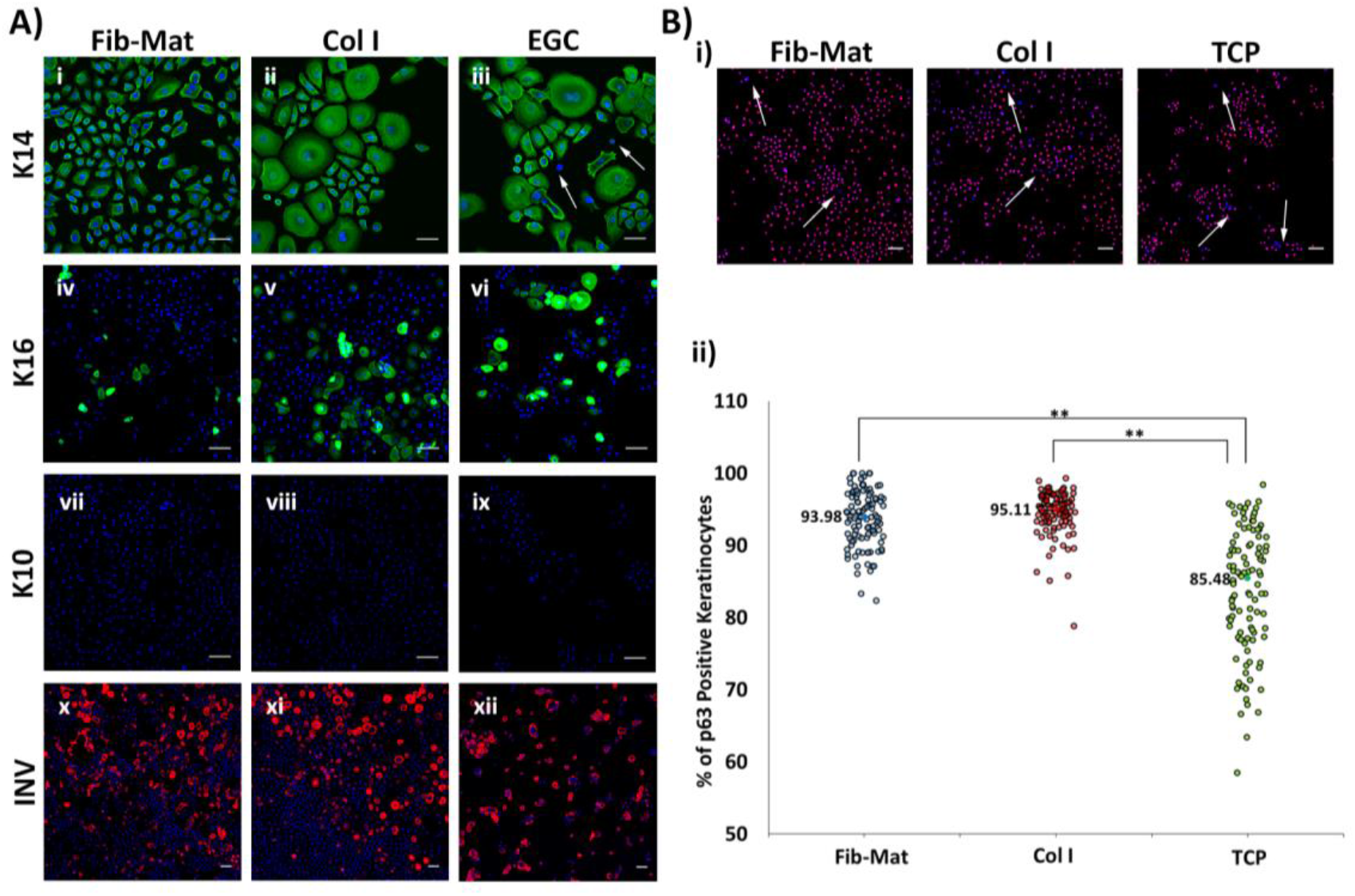
Expression of differentiation markers by keratinocytes grown on different substrates. **A)** The expression of K14 (i-iii), K16 (iv-vi), K10 (vii-ix) and involucrin (x-xii) by keratinocytes grown for three days on Fib-Mat, Col I and etched glass coverslips (ECG). Nuclei were stained using DAPI (Blue). Scale bars are 50 μm for K14 and 100 μm for K10, K16 and involucrin. Arrows indicate keratinocytes with no K14 expression. **B)** Expression of p63 by keratinocytes grown on Fib-Mat, Col I and TCP. i) Representative image of keratinocytes immunostained for p63. Arrows indicate the area of p63 negative keratinocytes. The nuclei were stained using DAPI (Blue) Scale bars are 100 μm. ii) Quantification of keratinocytes positive for p63. The data shown are representative of three separate experiments. Median values are given. Statistical analyses were a Kruskal-Wallis test followed by a Mann-Whitney test. ** = P<0.01

Keratinocytes on each of the substrates were also examined for p63 expression, a marker of “stemness”. As is shown in Figure 5B, while p63 expression was detected in keratinocytes on all substrates, the number of cells expressing p63 differed. The number of p63 expressing cells was significantly (p<0.01) higher on Fib-Mat (93.98%) and type I collagen (95.11%) than on TCP (85.48%) and the percentage of p63 positive cells was similar on both Fib-Mat and type I collagen (Fig. 5Bii).

Keratinocytes grown on Fib-Mat appeared to be smaller than keratinocytes similarly cultured on type I collagen or TCP (Fig. 6Ai). An assay was established based on Haase *et al*.[25], where cell size was determined by the area covered by the cell. Analysis of the size of individual keratinocytes revealed a statistically significant difference (p≤0.05) in the size of keratinocytes grown on Fib-Mat or type I collagen. The majority of keratinocytes residing on Fib-Mat were small cells, whilst type I collagen had the greatest number of large flat keratinocytes (Fig. 6B). While there were differences in size between cells on Fib-Mat and TCP, the differences were not statistically significant.

**Figure 6:**
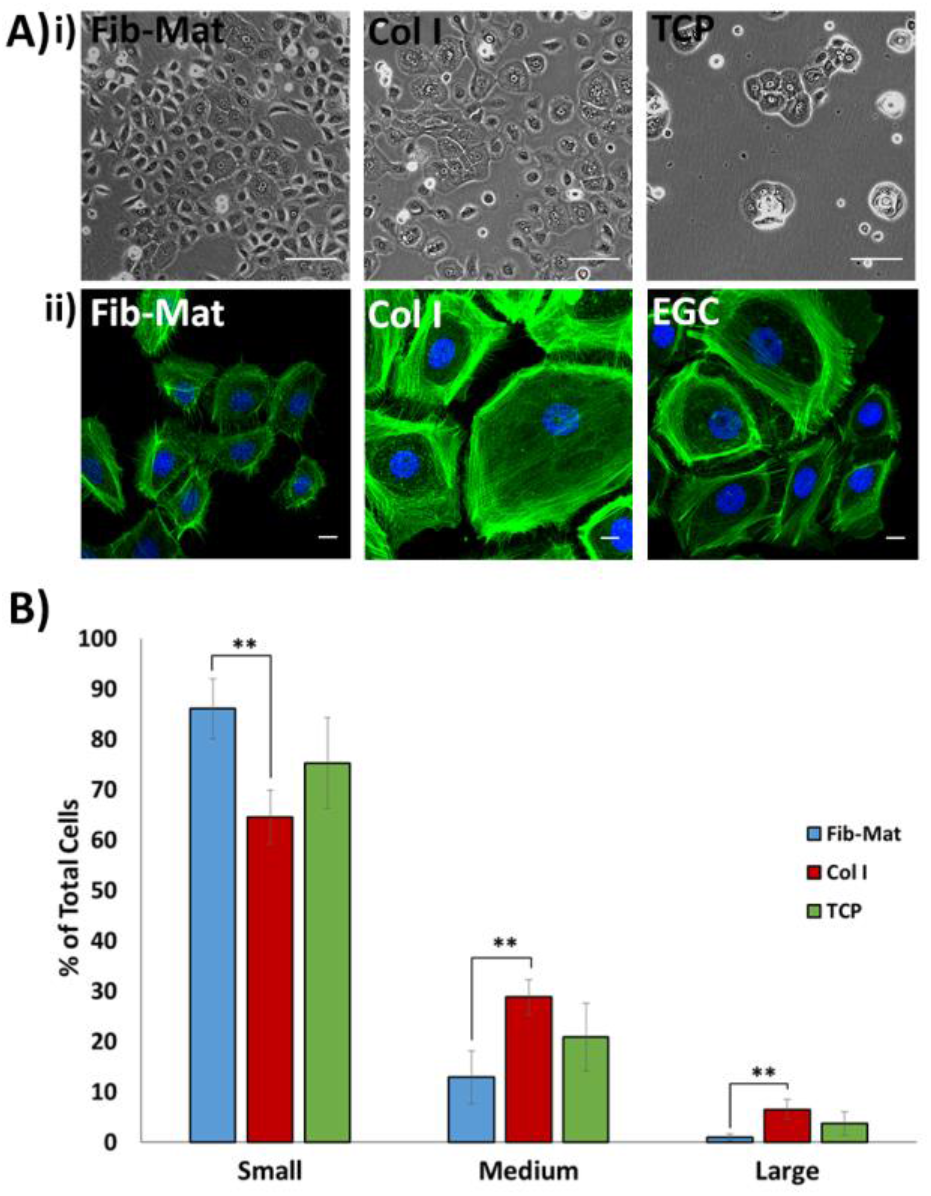
Size of keratinocytes grown on different substrates. **A)** i) Phase contrast images show differences in the size of keratinocytes grown on the various substrates. Images were captured on day 3 of culture. Scale bars are 100 μm. ii) Representative images of keratinocytes stained with phalloidin-Alexa Fluor^®^ 488 demonstrating that phalloidin staining accurately revealed keratinocyte size. Scale bars are 50 μm. Nuclei were stained using DAPI (Blue). **B)** Frequency of keratinocytes of differing size. Cell size was categorised as small, medium or large based on cell area (small<medium<large = <2000μm^2^<4000μm^2^<6000μm^2^). The data shown are the mean percent of total cells ± standard deviation. Pooled data from three separate experiments are shown. *=P≤0.05.

To evaluate the self-renewal capability of keratinocytes grown on different substrates, their colony forming ability was examined. Keratinocytes grown on either Fib-Mat, type I collagen or TCP were harvested, then seeded onto a layer of mitomycin-c treated feeder cells and grown for 12 days. The number of large colonies produced by keratinocytes grown on Fib-Mat was significantly higher (p<0.01) than that seen for keratinocytes from either type I collagen or TCP. Furthermore, more colonies regardless of size were observed in cultures of keratinocytes from the Fib-Mat (Fig. 7).

**Figure 7:**
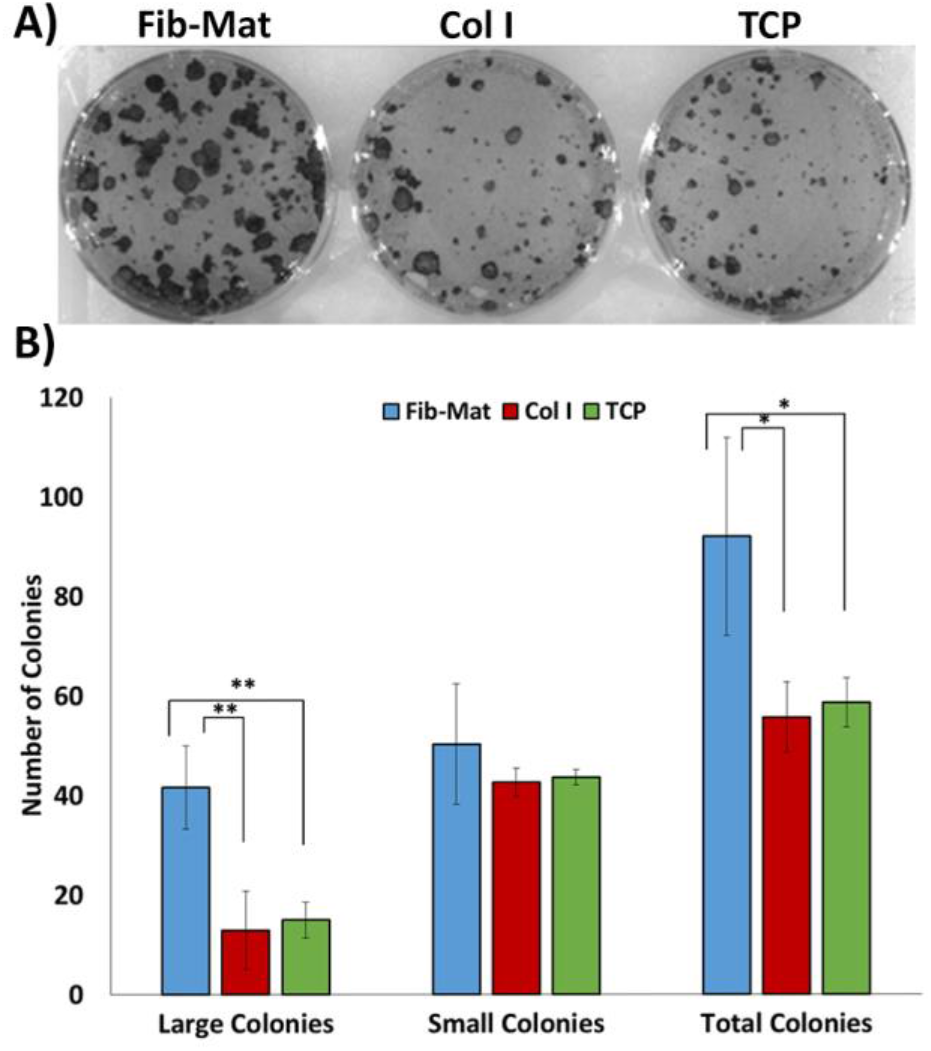
Colony forming ability of keratinocytes previously cultured on different substrates. **A)** Representative image of the colonies formed by keratinocytes previously cultured on different substrates. Keratinocytes were cultured for three days on Fib-Mat, type I collagen or uncoated TCP, the cells were harvested and single cells were seeded at low density onto mitomycin treated 3T3-J2 feeder cells. After 12 days keratinocyte colonies were stained with toluidine blue and imaged. **B)** Quantification of the colonies. All colonies were counted and categoried: large colonies were ≥ 1mm^2^, and small colonies were cell clusters <1mm^2^. The data shown are representative of three separate experiments. * = P≤0.05 and ** =P<0.01

### Keratinocytes are Highly Motile on Dermal Fibroblast-Derived Matrix

Cells were seeded onto EGC or TCP either uncoated, or coated with Fib-Mat or type I collagen, and left overnight to adhere before time-lapse images were taken at 15-minute intervals over a 2 day period. Distinct keratinocyte colonies were only seen when cells were on uncoated EGC or TCP. Pseudo-colonies, where keratinocytes migrated as a group of cells to form a colony, but then dispersed or combined with other colonies, were observed for cells on Fib-Mat regardless of the underlying surface. Keratinocytes on type I collagen coated EGC also formed pseudo-colonies, whereas on TCP coated with type I collagen, keratinocytes migrated as single cells. The majority of keratinocytes grown on either surface coated with Fib-Mat were highly motile over the entire 2 day period. Initially keratinocytes were similarly motile on type I collagen coated EGC or TCP, but as time progressed a proportion became less motile, and reduced motility was accompanied by an increase in cell size. Keratinocytes on uncoated EGC or TCP were the least motile, and an increase in cell size was also observed. (Links to videos are in supplementary information.)

Cell motility is linked to the organization of the actin cytoskeleton. To examine the arrangement of filamentous actin (F-actin) keratinocytes were grown on Fib-Mat, type I collagen or EGC for 3 days before being stained with phalloidin-Alexa Fluor^®^ 488. On type I collagen and EGC, well-developed actin stress fibres were observed at the keratinocyte circumference. This was more prominent in the large keratinocytes. In contrast, stress fibres at the cell circumference were less visible in keratinocytes on Fib-Mat (Fig. 6Aii).

## Discussion

The regenerative ability of keratinocyte stem cells has been known since the 1980s and has been well described in numerous studies[26–30]. However, the expansion of keratinocytes *in vitro* for clinical use has remained challenging, at it is dependent on harvesting a sufficient numbers of keratinocyte stem cells and their survival during propagation. The traditional “Rheinwald and Green method” uses murine feeder cells and FBS[4] and although this system has been used successfully it will face future hurdles as regulators embrace xenogeneic free systems as the norm for producing cells for clinical use. Keratinocytes can be expanded *in vitro* using a defined serum-free medium, and a collagen matrix to support cell attachment and growth[6, 7, 31], however, prolonged culture of keratinocytes in this way induces phenotypic changes; specifically a diminished capacity for self-renewal and an increased commitment towards terminal differentiation or senescence[7, 32, 33]. A critical factor for the long-term expansion of keratinocyte stem cells is their natural microenvironmental niche[8] and a key component of this niche that is lacking in the current defined culture system is a native ECM. In an effort to recapitulate a similar niche that the keratinocytes inhabit *in vivo*, we developed a method to generate a xenogeneic free dermal Fib-Mat (Fig. 1). This Fib-Mat substrate and defined serum-free medium better supported the proliferation of undifferentiated keratinocytes, compared to cultures of the same keratinocytes in defined serum-free medium on substrates of type I collagen or tissue TCP.

While the production and use of cell derived matrices to support the proliferation of undifferentiated stem cells has been employed previously [14, 16, 34–37], the generation of xenogeneic-free Fib-Mat using a tissue culture process that includes macromolecular crowding during the deposition of the ECM, and PLA_2_ for decellularisation, is novel. The inclusion of macromolecular crowding reagents in the primary dermal fibroblast xenogeneic-free cultures was found to reproducibly produce matrices that completely covered the surface of the culture plates. The PLA_2_ decellularisation protocol generated the most acceptable acellular matrices in terms of absence of immunological relevant remnants, and ECM structure preservation. Proteomics analysis of Fib-Mat generated under PLA_2_ had a core matrisome protein composition that was similar to that reported for the dermis [38, 39]. Interestingly, compared to proteomic data derived from skin dermis, the Fib-Mat lacked many of the matrisome-associated proteins. As some of the matrisome-associated proteins actually are cell-associated proteins (e.g. LGALS1, LGALS3 & GPC1) present in a full skin biopsy, it would be therefore be absent in the decellularised Fib-Mat as the cells were removed. Furthermore, as Fib-Mat is the result of a monolayer culture of fibroblasts, ECM proteins contributed by other residential cells (e.g keratinocyte & melanocyte) within the skin dermis will also be absence.

The ability of Fib-Mat to support proliferation of keratinocytes in the serum-free medium, DKSFM, was compared to similar cultures on type I collagen, the recommended substrate for use with DKSFM for keratinocyte propagation[6, 7] with TCP and no protein coating being the negative control. Keratinocytes adhered better to Fib-Mat compared to their adhesion on type I collagen or TCP. Greater keratinocyte proliferation as determined by cell number and Ki67 expression occurred on Fib-Mat, compared to keratinocytes on the other substrates. These data agree with other studies using stem cells. These other studies reported that cell-derived matrices which matched the tissue microenvironment of the cells *in vivo* better supported the attachment and proliferation of mesenchymal stem cells[40–42] and synovium-derived stem cells [43, 44]. We observed an initial lag in the proliferation of keratinocytes on Fib-Mat, before an exponential increase in proliferation occurred. As Fib-Mat is a complex biological substrate compared to type I collagen, the acclimatisation of keratinocytes towards this substrate may explain the initial slow proliferation rate.

Keratins are expressed by keratinocytes in a site-specific and differentiation-dependent manner. *In vivo* K14 is expressed in keratinocytes in the basal layer of the epidermis. However, as keratinocytes differentiate and migrate towards the surface of the epidermis, K14 expression is downregulated[45, 46]. Studies assessing the *in vitro* culture of keratinocytes in which K14 has been knocked down, showed decreased K14 expression was associated with reduced cell proliferation and an increase in the differentiation markers K10 and involucrin[47, 48]. In our study, K14 was expressed by keratinocytes grown on all three substrates, however there were a small proportion of keratinocytes on EGC without K14 staining (Fig. 5A). No K10 expression was observed regardless of the substrate the keratinocytes were cultured on. In contrast, involucrin was expressed by keratinocytes grown on all substrates, with a higher proportion of keratinocytes being involucrin-positive on uncoated EGC. The loss of K14 and the increased involucrin expression coincided with decreased keratinocyte proliferation on TCP/EGC. These data suggest that keratinocytes on TCP/EGC are more prone to initiating differentiation in the absence of stratification, whereas the continued K14 expression by keratinocytes residing on Fib-Mat is consistent with their preserved growth potential. However, K14 expression was not tightly associated with cell proliferation rates because there was no apparent decrease in K14 expression, despite reduced keratinocyte proliferation on type I collagen (Fig 4-5).

The pattern of K16 expression was informative, as very few keratinocytes grown on the Fib-Mat expressed this keratin whereas on the other substrates a high proportion of cells were K16 positive. K16 expression is associated with keratinocyte proliferation and migration and this keratin has been described as a marker of “stressed” or activated keratinocytes[49]. Keratin 16 is commonly expressed by keratinocytes in hyperproliferative diseases like psoriasis, squamous carcinoma and in wound healing. Indeed, within 6 h of wounding epithelial cells at the wound edge upregulate K16 expression and down regulate K10 expression[50] and these K16 expressing keratinocytes are involved in re-epithelization of wound site. From our data it is clear that K16 expression can be uncoupled from proliferation. Moreover, the low level of K16 expression by keratinocytes on Fib-Mat indicates that despite their increased proliferation these cells are not mimicking a wound healing response.

A functional link between p63 expression and keratinocyte stem cell maintenance within the skin has been shown by Mills *et al*.[51] and Yang *et al*.[52] using a p63−/− mouse. Further studies by Parsa *et al*.[53] showed that p63 expression is restricted to keratinocytes with high proliferative potential that reside within the basal layer. They also found that p63 is absent from terminally differentiating keratinocytes. Our study found most keratinocytes grown on Fib-Mat or type I collagen expressed p63, whereas, fewer keratinocytes expressed p63 when cultured on TCP (Fig. 5B). Interestingly, p63 expression coincided with a decline in growth potential (Fig. 4C) and an increase in involucrin expression (Fig. 5A). This inability of TCP to support the growth of undifferentiated keratinocytes is consistent with the data of others[6].

Numerous investigator have described cell size as a criteria distinguishing keratinocyte stem cells from keratinocytes committed towards differentiation[8, 54, 55]. Moreover, increased size is associated with differentiation, a characteristic that has been observed *in vivo* during normal epithelial maturation and in *in vitro* cultures [7]. Hence, the change in keratinocyte size, may indicate keratinocytes undergoing terminal differentiation, as keratinocyte enlargement accompanied by the expression of involucrin has been reported[56, 57]. Other studies also found that small keratinocytes are undifferentiated and retain a high proliferative capability[54, 55, 58]. We found more small keratinocytes were present in cultures grown on Fib-Mat (85%), whereas culturing the same keratinocyte population on type I collagen produced larger cells (Fig. 6). These data were consistent with what Esteban-Vives *et al*.[7] reported.

Recently Nanba *et al*.[59] suggested that cell motility is an attribute of undifferentiated keratinocytes. They found keratinocyte colonies with a high rotational movement, and in which individual cells were very motile, was indicative of these colonies containing undifferentiated keratinocytes with high proliferative capability. We found keratinocytes grown on Fib-Mat to be very motile (See video in supplementary information). Distinct keratinocyte colonies formed only on TCP, whereas pseudocolonies formed on Fib-Mat and type I collagen. Most keratinocytes grown on Fib-Mat were highly motile throughout the experiment, but keratinocytes cultured on type I collagen and TCP were less motile, a trait very evident when the time in culture was extended. Moreover, the reduced motility of individual keratinocytes was accompanied by an increase in cell size. The association of increased size, with decreased motility during keratinocyte differentiation was reported many years ago by Sun *et al*.[56] and our results are consistent with these findings.

Actin filament reorganisation is essential for changes in cell shape and motility. In our study keratinocytes plated on type I collagen or EGC without a matrix protein coating developed a circumferential actin network (Fig. 6Aii), similar to that reported by Nanba *et al*.[60] and which was described to be indicative of reduced cell movement and terminal differentiation. In contrast, keratinocytes grown on Fib-Mat had short bundles of actin that were radially distributed (Fig. 6Aii), an arrangement of actin filaments described as indicative of proliferative, undifferentiated keratinocytes[60].

Collectively, our data indicate that keratinocytes grown on Fib-Mat are less differentiated than keratinocytes cultured on TCP, which show signs of undergoing early commitment to terminal differentiation. Whereas, despite exhibiting characteristics that are indicative of differentiation (e.g. cell size, cell motility, lower colony forming ability and actin reorganization) keratinocytes grown on type I collagen still expressed markers (K14 and p63) of undifferentiated keratinocytes. Others have also found that keratinocytes retain some markers characteristic of undifferentiated cells during the early stages of differentiation. Webb *et al*.[61] found that keratinocytes in the basal layer of the epidermis do not switch off the expression of keratin 15 (a marker of keratinocyte quiescence, and in some circumstances of stem cells) even during the differentiation process. Furthermore, Esteban-Vives *et al*.[7], observed that keratinocytes grown on type I collagen still retained K15 expression despite showing signs of differentiation. Hence, it is likely that on type I collagen, the keratinocytes are in the early stages of terminal differentiation, even though some markers of undifferentiated cells are present.

This conclusion was further supported by the behaviour of the keratinocytes expanded on different substrates in a colony forming assay. Barrandon *et al*.[62] described the use of colony forming assays as an invaluable tool for determining the presence of stem cells within a keratinocyte population. Undifferentiated keratinocyte stem cells are described to have a higher self-renewal capability and form large, progressively growing colonies (> 1 mm^2^; holoclones). We found that keratinocytes grown on Fib-Mat produced a higher number of large colonies, when compared to keratinocytes expanded on the other two substrates. This indicate that Fib-Mat better retained and promoted the self-renewal ability of undifferentiated keratinocyte stem cells.

Others have also shown cell-ECM interactions are important for preserving the self-renewal ability of cultured keratinocytes. Adams and Watt[63] demonstrated that keratinocytes losing ECM contact are triggered to terminal differentiation. Similarly, our data and that of Coolen *et al*.[6], showed that keratinocytes undergo terminal differentiation when grown on tissue culture plastic that lacked an ECM protein. Hence, ECM proteins such as type I collagen[7], type IV collagen[6] and fibronectin[64] have been used as substrates to culture keratinocytes. Although using these single ECM proteins enable the keratinocytes to adhere and proliferate, they do not sustain the long-term growth of keratinocytes[7, 32]. In this reductionist approach, the synergistic impact of growth factors and ECM proteins and their coordinated signalling pathways in the keratinocytes is overlooked. Others have shown that even the combination of three matrix proteins can have a synergistic effect[65–67]. Flaim *et al*.[65] found that the combination of type I collagen with laminin and type III collagen enabled embryonic stem (ES) cells to efficiently differentiate towards a liver progenitor lineage, although individually these matrix proteins were unable to promote liver progenitor cell differentiation. Furthermore, Watt *et* al.[64] showed that substrates comprising a combination of laminin, type IV collagen and fibronectin inhibited the differentiation of keratinocytes during *in vitro* culture.

The proteomics data indicated that our Fib-Mat contained laminins, type IV collagen and fibronectin plus numerous other ECM proteins, and some, but not all, the ECM associated proteins, but very few of the secreted factors found in the dermis. Our data indicate the combined signals of the core matrisome proteins that are present in Fib-Mat, are sufficient to suppress keratinocyte differentiation and to promote proliferation. It is not possible to say from our data that the complex mixture of chemokines, growth factors and other secreted factors are also contributing to suppressing keratinocyte differentiation. In our study the essential growth factors for keratinocyte proliferation were probably present in the culture medium and the Fib-Mat provided the ECM components to correctly present these growth factors to the growing keratinocytes. However, whether essential secreted factors were present in the Fib-Mat at very low concentration is unclear as the proteomics methodology may not detect such proteins.

In conclusion, this study highlights the important role of a native ECM in modulating keratinocyte growth and differentiation. Our novel culture system using dermal fibroblast ECM is superior to current protocols for the serum-free culture of keratinocytes for clinical use because it delivers undifferentiated keratinocytes that continue to proliferate. In contrast, keratinocytes expanded using type I collagen (the current protocol) are likely to have progressed down the terminal differentiation pathway as a result of the expansion protocol used.

## Acknowledgements

The authors are grateful to Dr. Paula Benny for her technical assistance and Mr. John Lim for assisting in image analysis. The authors would also like to thank A/Prof. Pritinder Kaur for her guidance in the colony formation assay and Prof. John Whitelock and Dr Megan Lord for their gift of the anti-perlecan antibody, CCN-1. Chee Wai Wong was supported by a Curtin University Strategic International Research Scholarship. Radoslaw M Sobota was funded by IMCB, A*STAR, Young Investigator Grant YIG 2015 (BMRC, A*STAR) and NMRC MS-CETSA platform grant MOHIAFCAT2/004/2015. The authors acknowledge the provision of research facilities and technical assistance of the staff of CHIRI Bioscience, which was partly funded by Curtin University, State and Commonwealth Governments.

## Author Contribution

Chee-Wai Wong: Conception and design, collection and assembly of data, data analysis and interpretation, manuscript writing, final approval of manuscript.

Beverley F. Kinnear: Conception and design, final approval of manuscript.

Radoslaw M. Sobota: Conception and design, collection and assembly of data, final approval of manuscript.

Rajkumar Ramalingam: Conception and design, collection and assembly of data, final approval of manuscript.

Catherine F. Legrand: Collection and assembly of data, final approval of manuscript. Danielle E. Dye: Conception and design, final approval of manuscript.

Michael Raghunath: Conception and design, final approval of manuscript.

E. Birgitte Lane: Conception and design, supply of research materials, financial support, final approval of manuscript.

Deirdre R. Coombe: Conception and design, data interpretation, financial support, manuscript writing, final approval of manuscript.

---

**Figure S1:**
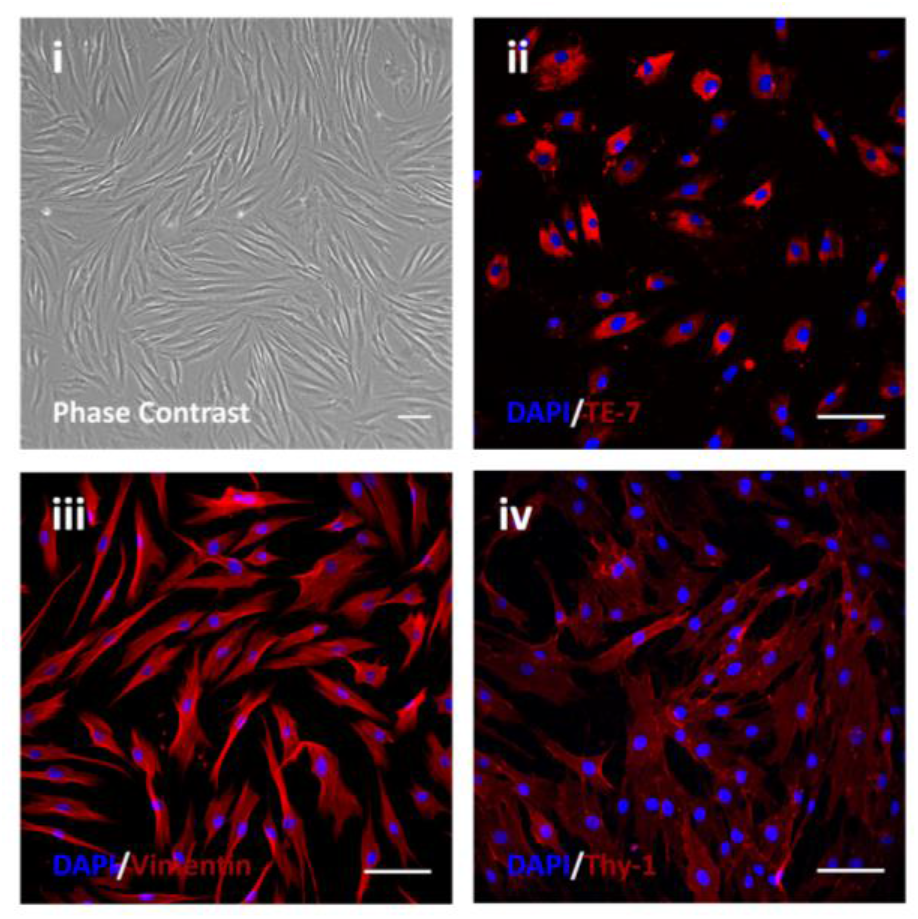
Characterization of human dermal fibroblasts (HDF). HDF were stained with antibodies recognising the fibroblast markers: TE-7 (ii), vimentin (iii) or Thy-1 (iv). The secondary antibody was an Alexa Fluor^®^ 546-conjugated anti-mouse IgG1. Nuclei were stained with DAPI (blue). Scale bars are 100 μm.

**Figure S2:**
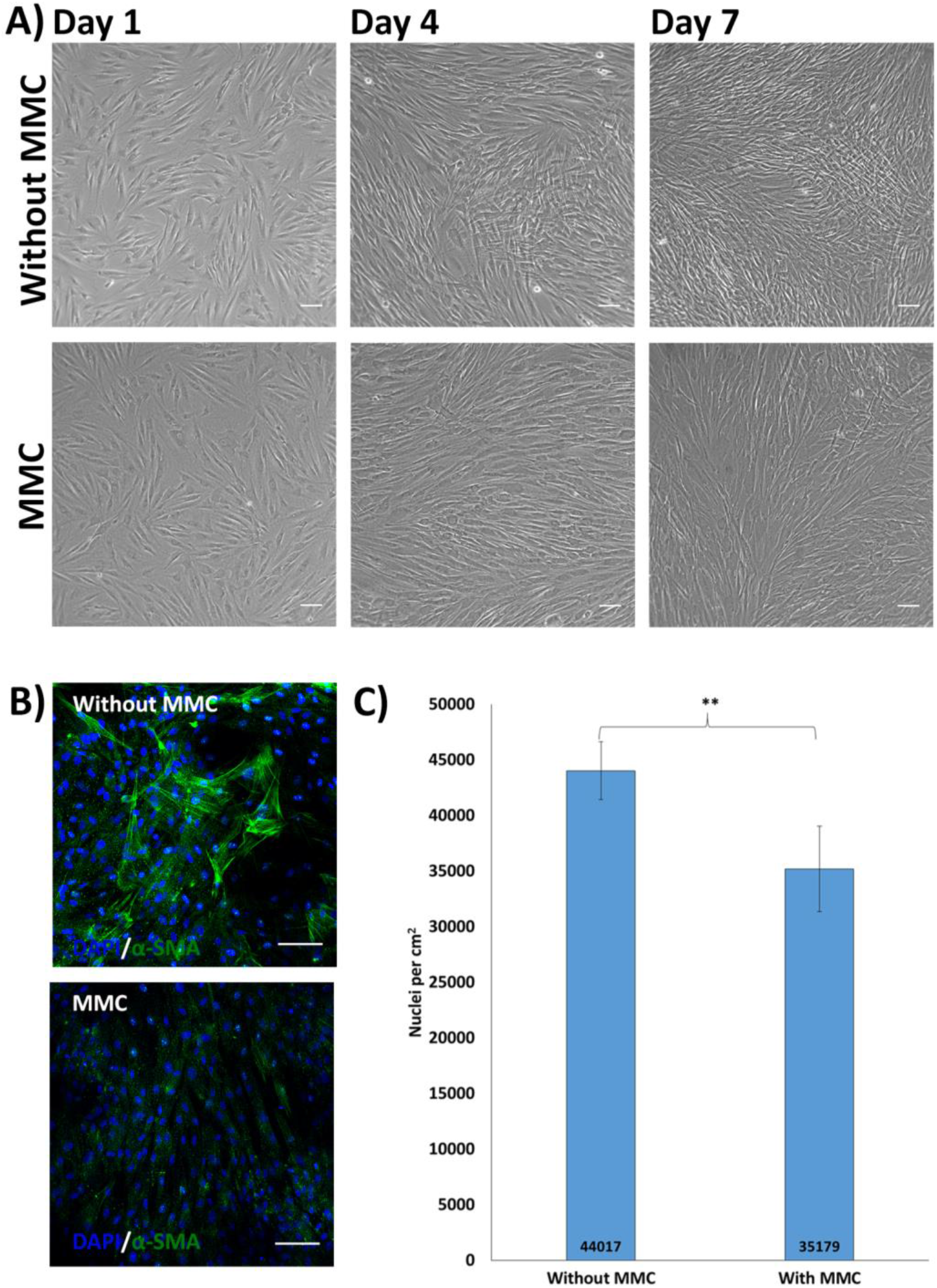
The effect of MMC on fibroblast behaviour. **A)** Phase contrast images of HDF cultured with or without MMC. Scale bars are 100 μm. **B)** Myofibroblast differentiation with and without MMC. HDF were grown either with or without MMC for seven days. The cells were stained with a mAb recognising α-smooth muscle actin (α-SMA) and imaged by confocal microscopy. The secondary antibody was an Alexa Fluor^®^ 488-conjugated anti-mouse IgG2a. Nuclei were stained using DAPI (Blue). Scale bars are 100 μm. **C)** HDF grow more slowly under MMC conditions. The number of HDFs after culturing with or without MMC for seven days. Data are expressed as mean ± SD. Mean values are given. Shown is a representative of triplicate experiments. ** = P <0.01.

**Figure S3:**
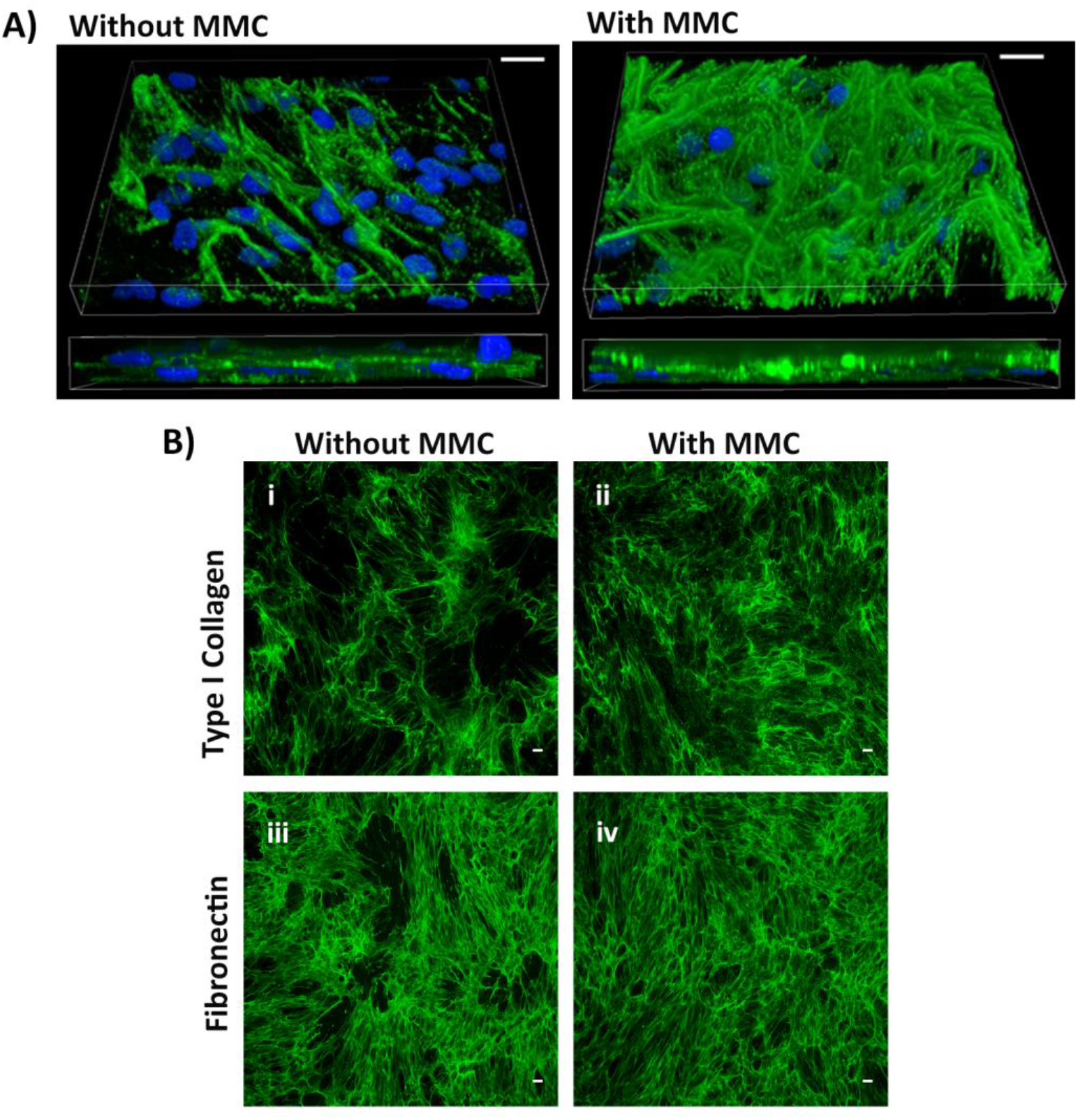
The effect of MMC on the ECM deposited by HDF. **A)** The effect of MMC on the 3D architecture of the ECM deposited by HDF. A 3D Z-stacked confocal image of type I collagen deposited by HDFs grown without or with MMC for seven days. Cells and matrix were immunostained for type I collagen. The secondary antibody was an anti-rabbit IgG Alexa Fluor^®^ 488-conjugated antibody. Nuclei were stained with DAPI (Blue). Scale bars are 10 μm. **B)** The ECM deposited by HDF with or without MMC after decellularisation. The HDF were grown with or without MMC for seven days. The cell layers were decellularised using PLA_2_. Matrices were immunostained for type I collagen (i, ii) or fibronectin (iii, iv). The secondary antibody was an Alexa Fluor^®^ 488-conjugated anti-rabbit IgG. Nuclei were stained with DAPI (Blue). Scale bars are 100 μm.

**Figure S4:**
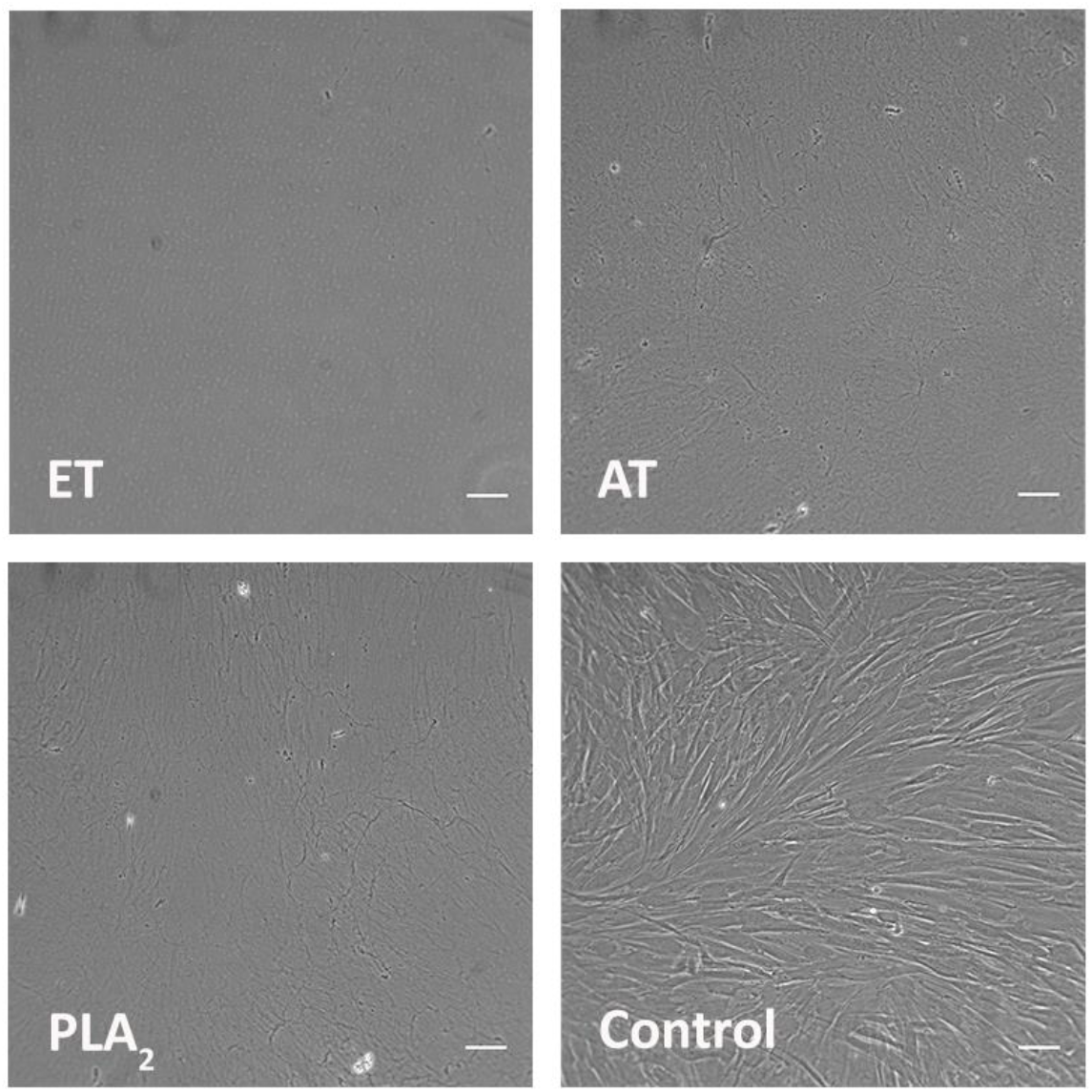
Decellularised dermal fibroblast-derived ECM prepared using EDTA, ammonia hydroxide (AH) or phospholipase A_2_ (PLA_2_). Images were obtained using phase contrast microscopy. The HDF were grown with MMC for seven days and decellularised as indicated. The control was HDF before decellularisation. Scale bars are 100 μm.

**Figure S5:**
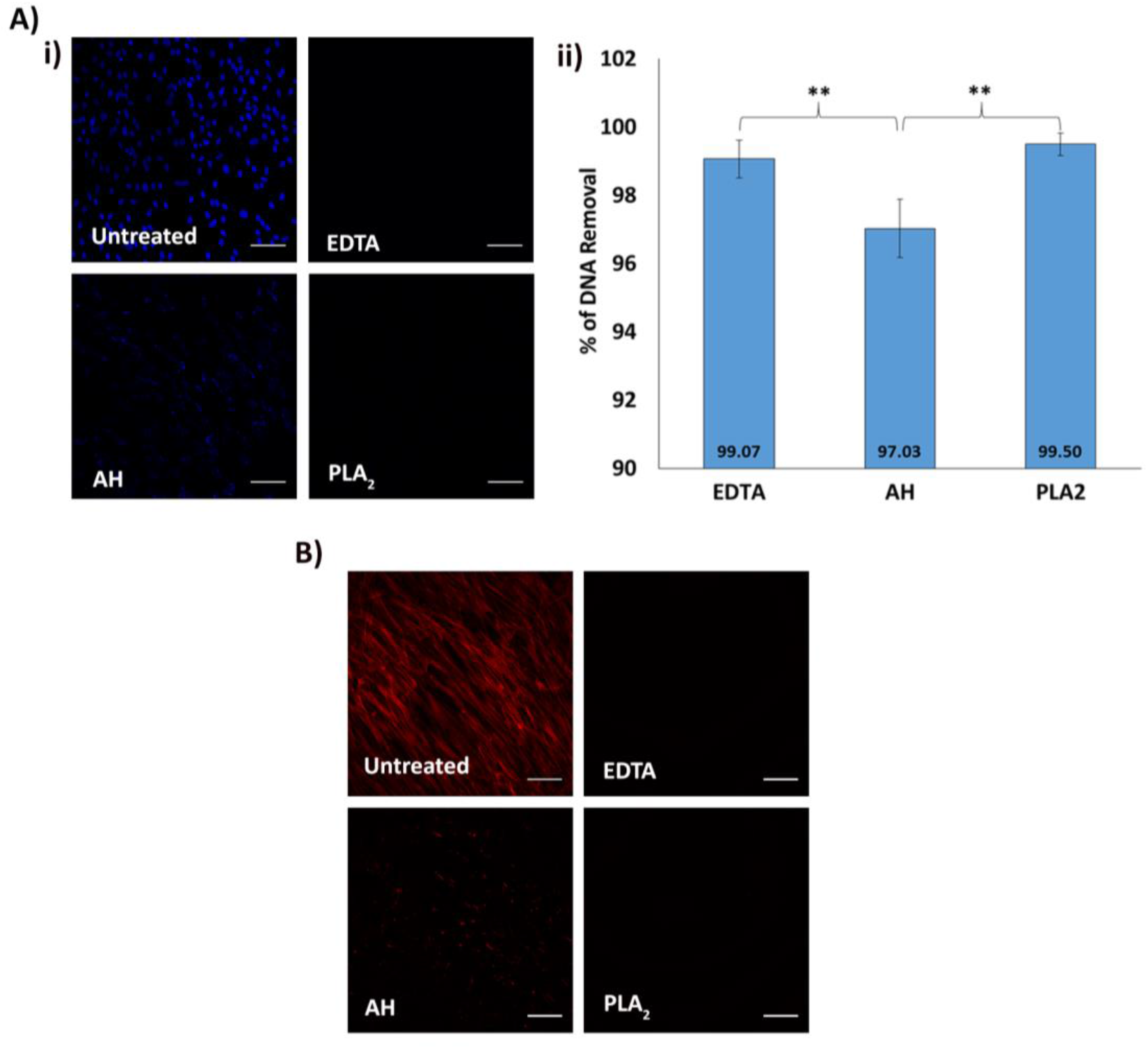
Efficacy of methods for decellularisation of HDF matrices. **A)** Efficacy of decellularisation methods for removing nuclear components. i) DAPI staining of the variously decellularised ECM and untreated HDF control. HDF were grown with MMC for seven days then decellularised by the method indicated. Images were obtained using fluorescence microscopy. Scale bars are 100 μm. ii) Quantification of DNA removed after decellularisation. The CyQuant dye was used to measure the DNA present following decellularisation and these fluorescent intensity values were subtracted from the fluorescent intensity of the untreated control to allow calculation of percent of DNA removed. Means ± SD of 4 replicates are shown. The figure is a representative of three experiments. ** = P<0.01. **B)** Efficacy of different decellularisation treatments for removing cytoskeletal components. Phalloidin staining of the variously decellularised ECM and untreated HDF control cell layer for polymerised actin. Images were obtained using fluorescence microscopy. Scale bars are 100 μm.

**Figure S6:**
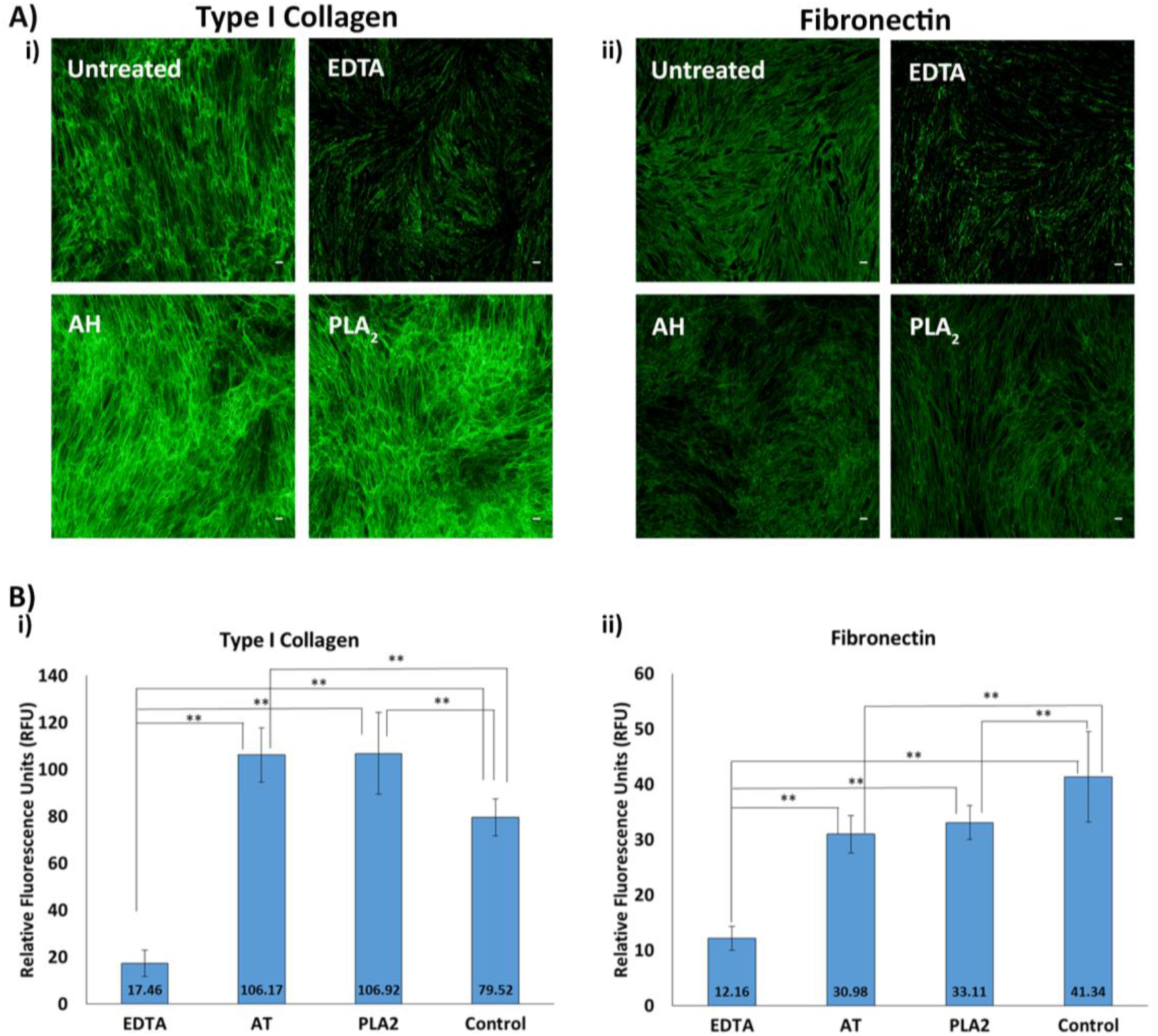
Quantities of the deposited ECM after decellularisation. **A)** Representative images of ECM obtained using different decellularisation treatments: EDTA, ammonia hydroxide (AH) & phospholipase A_2_ (PLA_2_). The acellular ECM were immunostained using antibodies recognizing either type I collagen or fibronectin. Scale bars are 100 μm. **B)** Quantification of fluorescence intensity of type I collagen (i) and fibronectin (ii) immunostaining after decellularisation. Data are expressed as mean ± SD. Mean value are shown. The figure is representative of three experiments. *=P<0.05

**Figure S7:**
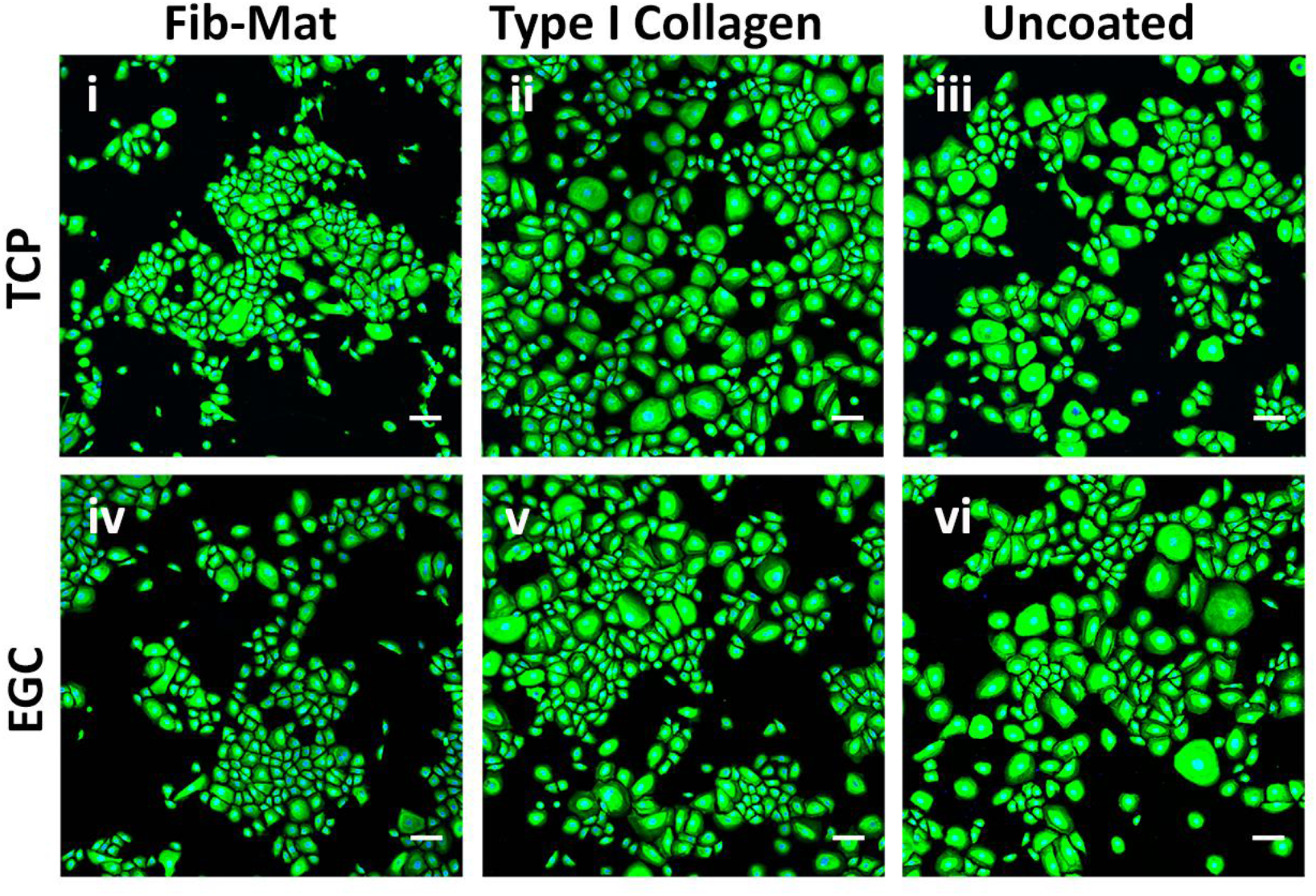
Comparison of tissue culture plastic (TCP) and etched glass coverslips (EGC) as a platform for keratinocyte growth. Keratinocytes grown on either EGC or TCP coated with either Fib-Mat (i,iv), type I collagen (3ug/cm2: ii,iv) or were uncoated (plain: iii, vi). Keratinocytes were cultured for 3 days in DKSFM then fixed with acetone: methanol (1:1) and immunostained for K14. The secondary antibody was an anti-mouse IgG Alexa Fluor^®^ 488-conjugated antibody (Green). Nuclei were stained with DAPI (Blue). Scale bars are 100μm.

